# Scelestial: fast and accurate single-cell lineage tree inference based on a Steiner tree approximation algorithm

**DOI:** 10.1101/2021.05.24.445405

**Authors:** M.-H. Foroughmand-Araabi, S. Goliaei, A. C. McHardy

## Abstract

Single-cell genome sequencing provides a highly granular view of biological systems but is affected by high error rates, allelic amplification bias, and uneven genome coverage. This creates a need for data-specific computational methods, for purposes such as for cell lineage tree inference. The objective of cell lineage tree reconstruction is to infer the evolutionary process that generated a set of observed cell genomes. Lineage trees may enable a better understanding of tumor formation and growth, as well as of organ development for healthy body cells. We describe a method, Scelestial, for lineage tree reconstruction from single-cell data, which is based on an approximation algorithm for the Steiner tree problem and is a generalization of the neighbor-joining method. We adapt the algorithm to efficiently select a limited subset of potential sequences as internal nodes, in the presence of missing values, and to minimize cost by lineage tree-based missing value imputation. In a comparison against seven state-of-the-art single-cell lineage tree reconstruction algorithms - BitPhylogeny, OncoNEM, SCITE, SiFit, SASC, SCIPhI, and SiCloneFit - on simulated and real single-cell tumor samples, Scelestial performed best at reconstructing trees in terms of accuracy and run time. Scelestial has been implemented in C++. It is also available as an R package named RScelestial.

## 1 Introduction

Lineage trees describe the evolutionary process that created a sample of clonally related entities, such as individual cells within an organ or a tumor. A lineage tree also suggests the constitution of a founder cell, as well as of its descendants, and the evolutionary events that occurred during lineage formation. Single-cell genomic data provide a highly resolved view of cellular evolution, substantially more than bulk genome sequencing [1]. However, they also come with high rates of missing values and distorted allele frequencies arising from amplification biases and sequencing errors [2]. Lineage tree reconstruction from single-cell data therefore requires specific considerations for handling missing values and errors. Specific approaches to tackle this problem include the methods of Kim and Simon [3], BitPhylogeny [4], OncoNEM [5], SCITE [6], SiFit [7], SASC [8], SPhyR [9], SCIPhI [10], SiCloneFit [11], B-SCITE [12], and PhISCS [13]. Kim and Simon [3] make use of the infinite site assumption and infer a “mutation tree” based on calculating a probability for ordering mutations in a lineage and constructing a tree, finding the maximum spanning tree in this graph. BitPhylogeny [4] provides a stochastic process through a graphical model that stochastically generates the given input data and uses Markov Chain Monte Carlo (MCMC) for sampling to search for the best lineage tree model and the associated parameters. OncoNEM [5] and SCITE [6] infer a phylogenetic tree over all the samples under a maximum likelihood model. OncoNEM uses a heuristic search, whereas SCITE uses MCMC sampling to find a maximum likelihood tree. SiCloneFit [11] infers subclonal structures and a phylogeny via a Bayesian method under the finite site assumption. SASC [8] and SPhyR [9] consider the k-Dollo model, a more relaxed model in comparison than infinite site assumption. In k-Dollo model, a mutation can be gained once in a tumor but may be lost multiple times afterwards. SASC uses simulated annealing and SPhyR uses k-means to find the best k-Dollo evolutionary tree. B-SCITE [12] and PhISCS [13] infer subclonal evolution, based on a combination of single-cell and bulk sequencing data. B-SCITE uses an MCMC method to maximize a likelihood function for trees and sequencing data. PhISCS formulates tree reconstruction as combinatorial and mathematical programming problems, and uses standard mathematical programming solvers to find the solution. Integer or integer linear programmings for the phylogenetic tree reconstruction are also available [14, 15].

The Steiner tree problem is a classic problem in theoretical computer science with a wide range of applications in various areas including very-large-scale integration design [16], network routing [17], civil engineering [18], and other areas [19–22]. Given a weighted graph and some vertices marked as terminals, the Steiner tree problem is the problem of finding the minimum weighted tree that connects all the terminal vertices, with non-terminal vertices that may or may not be included in the optimal tree. Although the Steiner tree problem is NP-hard and no polynomial-time exact solution is known, some elegant approximation algorithms are available. They guarantee polynomial time and a constant factor approximation ratio, and have been used for phylogenetic reconstruction [23]. This is a great advantage compared with sampling heuristics such as MCMC for finding an optimal solution, which may be trapped in local optima.

Here, we describe Scelestial, a method for lineage tree reconstruction from single-cell datasets, based on the Berman approximation algorithm for the Steiner tree problem [24]. Our method infers the evolutionary history for single-cell data in the form of a lineage tree and imputes the missing values accordingly. Dealing with missing values makes the problem much harder in theory. To overcome this difficulty, we represent a hypercube represented via [24] as a sequence with missing values as its center. This representation, although imposes some cost in the results of the algorithm caused by misplacement of the center from the best imputation, which we do not know in advance, helps us to get a fast yet accurate and robust algorithm.

## 2 Results

### 2.1 Performance in lineage tree reconstruction on simulated data

We compared the performance of Scelestial to SCITE, OncoNEM, BitPhylogeny, SASC (as a recent instance of k-Dollo-based methods), SCIPhI, SiFit, and SiCloneFit. For this, we generated data with the cell evolution simulator provided by OncoNEM and with another tumor evolution simulator that we developed (Section 3.2). To assess the inferred trees, we calculated their distance to the ground truth lineage trees as the normalized pairwise distances between corresponding samples, as described in Section 3.4.1. In addition, we calculated the similarity between the generated trees and the ground truth by comparing tree splits (Section 3.4.2). For run time evaluation, we executed all the methods on simulated tumor data over a range of numbers of samples and sites. We also assessed the run time performance of the methods in relation to different parameters, such as the number of samples and sites (Section 2.5).

First, we evaluated the algorithms on data created with OncoNEM’s simulator. The simulator creates samples by producing clones and sampling from these clones (Section 3.3). It accepts the number of clones (i.e., the number of nodes in the lineage tree), the number of sampled cells, the number of sites, the false positive rate, the false negative rate, and the missing value rate as parameters. We used 1.5% as the setting for the false positive positives, 10% for false negatives and 7% for the missing value rate. These parameters are consistent with observed parameters in single-cell data [25]. We performed tests for 50 and 100 samples, 5 and 10 clones, and 20 and 50 sites, and calculated the pairwise distance between the inferred and ground truth trees for all methods (Table 1). From these data, Scelestial reconstructed lineage trees with the lowest distance to the ground truth tree in 5 out of 8 cases. SASC, SCITE, and SiFit each performed best in one of the remaining cases.

**Table 1:** Comparison of reconstruction of ground truth lineage tree from data simulated by OncoNEM, showing the distance between the inferred trees and the ground truth for all methods across eight lineage trees. The best results among all the methods for each evolutionary tree are shown in bold.

| Method | 50 samples |  |  |  | 100 samples |  |  |  |
| --- | --- | --- | --- | --- | --- | --- | --- | --- |
|  | 5 clones |  | 10 clones |  | 5 clones |  | 10 clones |  |
|  | 20 sites | 50 sites | 20 sites | 50 sites | 20 sites | 50 sites | 20 sites | 50 sites |
| BitPhylogeny | 0.8947 | 0.7949 | 0.8875 | 0.8665 | 1.056 | 0.9805 | 0.8418 | 1.1766 |
| OncoNEM | 0.9738 | 0.781 | 0.7412 | 0.8044 | 0.9783 | 0.9867 | 0.8429 | 0.9426 |
| SCITE | 1.0205 | 0.914 | 1.0122 | 1.2909 | 1.0366 | 0.8682 | 0.8536 | 1.0021 |
| SiFit | 0.9088 | 0.7383 | 0.7147 | 0.7543 | 0.9575 | **0.8603** | **0.784** | **0.861** |
| SASC | 0.8912 | 0.8212 | 0.7716 | 0.7994 | 0.9289 | 0.924 | 0.8186 | 0.9215 |
| SCIPhI | 0.9671 | 1.0897 | 0.7049 | 0.9466 | 1.0628 | 1.2485 | 1.2483 | 1.2194 |
| SiCloneFit | 1.0935 | 0.9441 | 0.8728 | 0.8433 | 1.0641 | 1.0464 | 0.809 | 0.9617 |
| Scelestial | **0.8904** | **0.703** | **0.6647** | **0.726** | **0.9241** | 0.8862 | 0.7918 | 0.8771 |

Next, we simulated 100 single-cell datasets from a solid tumor covering a range of evolutionary time spans (i.e., a range of mutations per branch in the resulting lineage trees), using a simulation method we implemented (Section 3.2, results in Fig. 1). The simulated data provide a granular simulation of tumor evolution and single-cell sequence data. Tumor growth was simulated with 50 samples and 200 sites. The other parameters (missing value rate, false positive rate, and false negative rate) were set as in the OncoNEM simulation. The main difference between our simulation and the OncoNEM simulation tool was that we considered the relative preference between clones.

**Fig 1:**
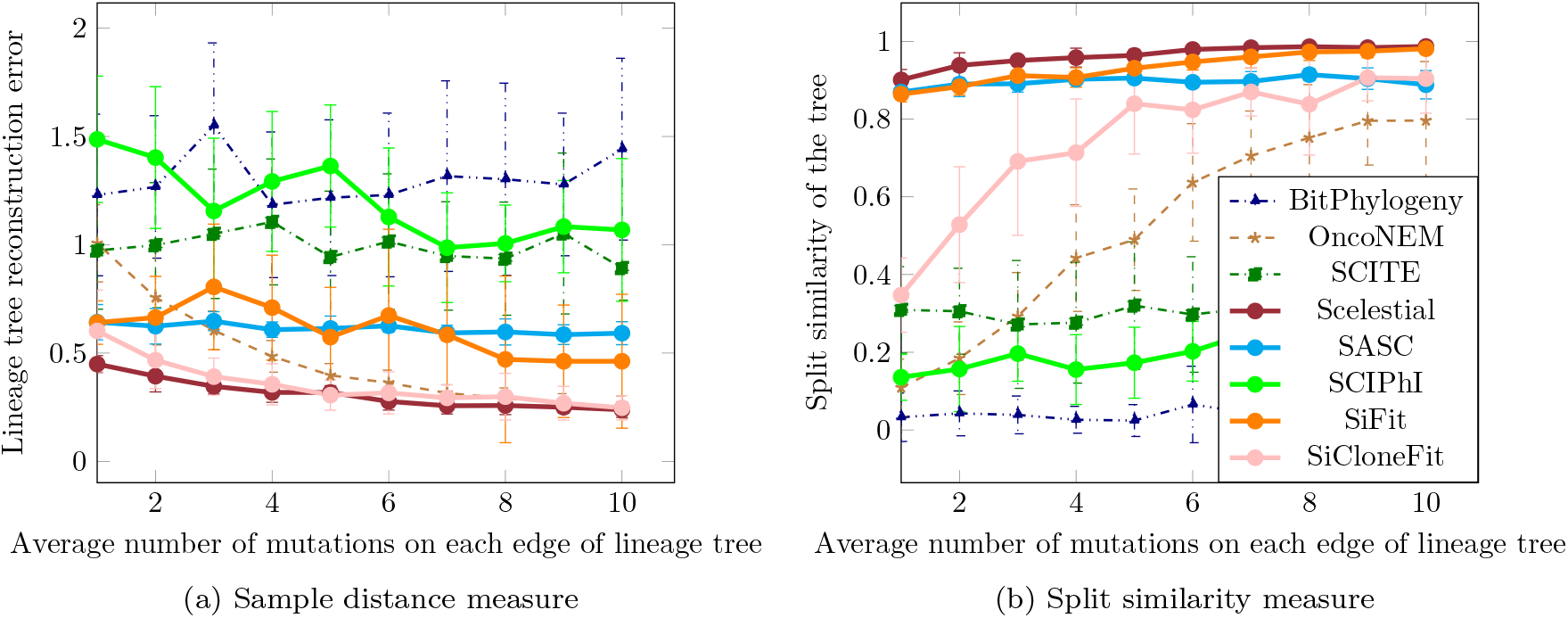
Comparison of the methods for single-cell lineage tree reconstruction on simulated tumor data. Note that in case of the lineage tree reconstruction error (Fig. 1a), lower values show a better reconstruction. On the other hand, the split similarity measure represents (Fig. 1b) similarity between the reconstructed tree and the ground truth tree, making higher values favorable.

The lineage tree reconstruction error was calculated as the sum of all distance errors over every pair of samples (Section 3.4.1). The lower the lineage tree reconstruction error, the more similar the tree is to the ground truth one. In addition, we determined the split similarity between the resulting trees and the ground truth tree (Section 3.4.2). A higher split similarity measure reflects a better reconstruction of the tree’s topology. Overall, when considering the lineage tree reconstruction error, none of the resulting trees was highly similar to the ground truth tree (Fig. 1a). Nevertheless, there were differences in the accuracy of the inferred trees. With these data, considering the average performance measure of 10 datasets for each mutation rate, Scelestial performed best in 10 out of 10 cases regarding the split similarity measure. Scelestial also performs best for the distance measure in 9 of 10 cases; SiCloneFit was the best for the remaining case. The performance of Scelestial, SiCloneFit, and OncoNEM increased with the number of mutations (Fig. 1a). This increase in the mutation rate had a very large effect on the performance of SiCloneFit and OncoNEM, whereas the performance of SASC and SiFit was almost stable across changes in the number of mutations between clones.

Overall, the relative performance of lineage tree reconstruction for different methods was similar to that observed before (Table 1). This also indicates that the relative difficulty of tree inference for the simulation method provided by OncoNEM and our simulation method are similar.

### 2.2 Robustness across varying data properties

We next evaluated the robustness of Scelestial to varying dataset properties over 420 datasets with varying parameters that we generated for this purpose with our tumor simulator. We fixed the default configuration in our simulation as 50 samples, 200 sites, 5 evolutionary nodes in the simulated evolutionary tree, 1.5% as the false positive rate, 10% as the false-negative rate, and 7% as the missing value rate. We set the average number of mutations between nodes in the evolutionary tree to be 20. For evaluating the robustness of Scelestial with respect to one parameter, we set the other parameters to their default values and evaluated Scelestial over a range of the parameter under study. We used the following ranges: false positive rate: 0–50%; false negative rate: 0–30%; missing value rate: 0–50%; samples: 5–200; sites: 20–1000.

Scelestial was not very sensitive to variation in the tested parameters (Fig. 2, Fig. 3) in terms of their influence on the tree distance error and topological accuracy measured by split similarity. As expected, performance decreased with increasing missing value, false negative, and false positive rates (Fig. 3). For datasets including more than 25 sites, the sample distance measure was stable.

**Fig 2:**
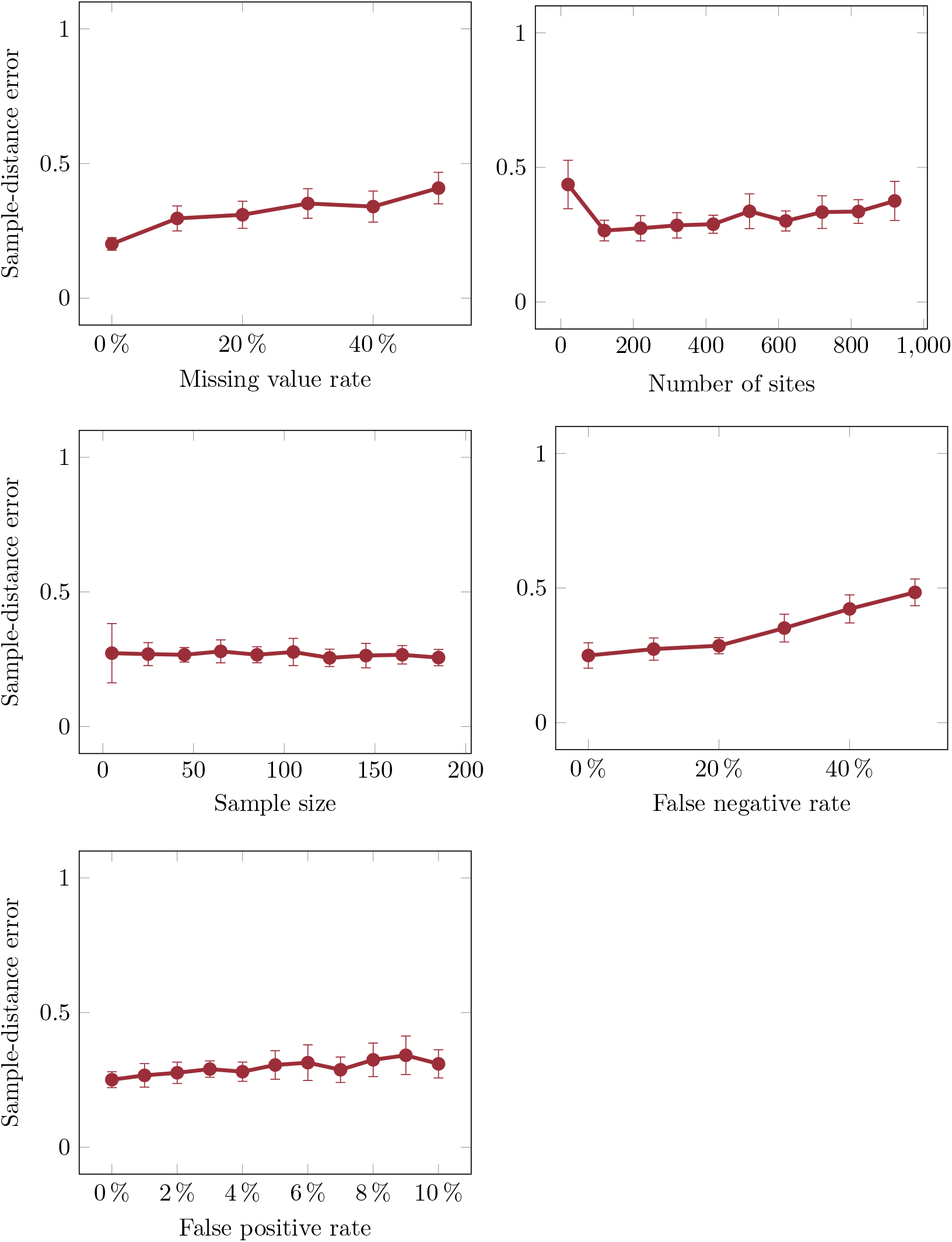
Robustness of Scelestial to variation in the properties of ground truth lineage trees in terms of sample distance in the trees between the inferred and ground truth trees.

**Fig 3:**
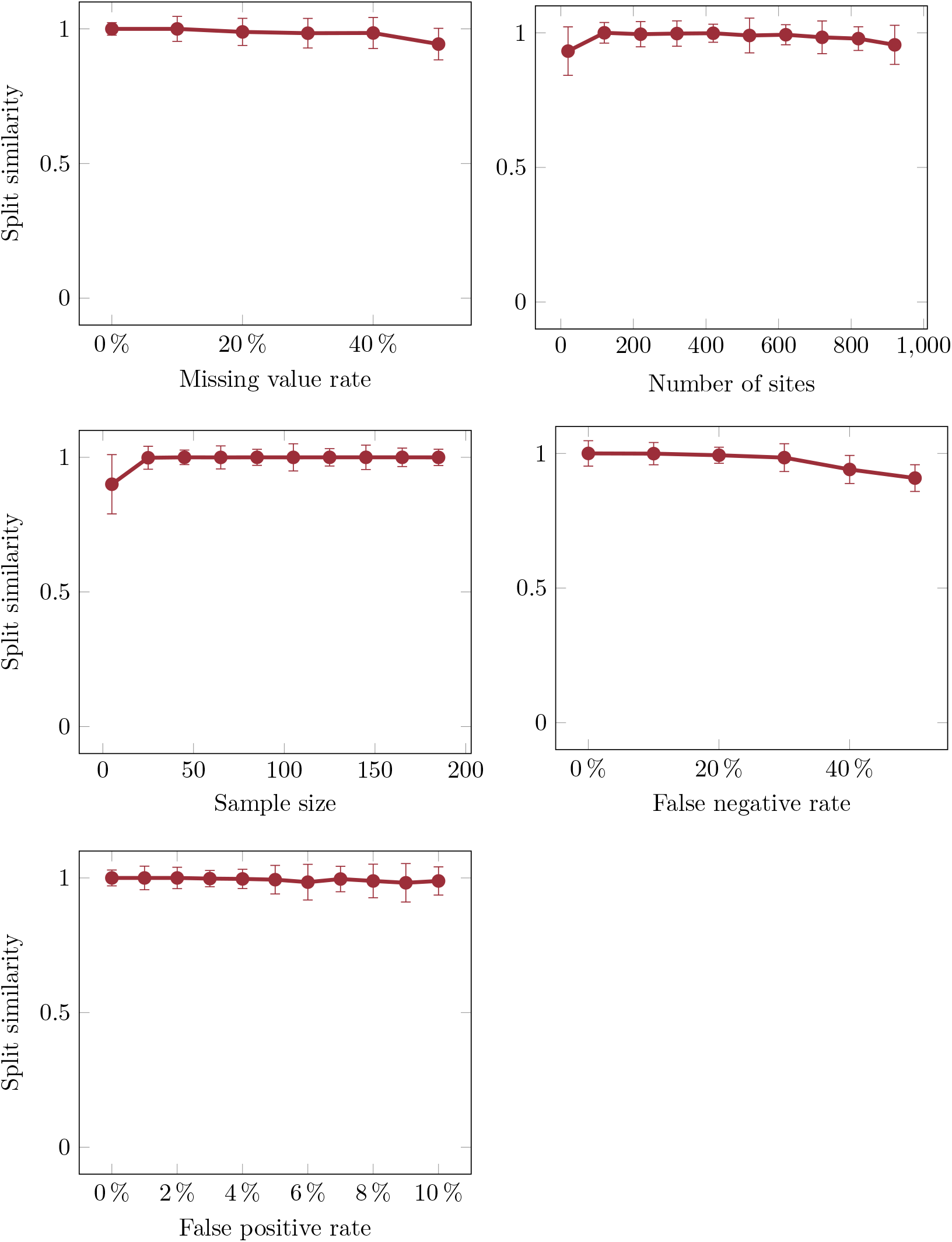
Robustness of Scelestial to variation in the properties of ground truth lineage trees in terms of topological similarity between the inferred and ground truth trees.

### 2.3 Case study: a single-cell dataset from a muscle-invasive bladder tumor

We inferred lineage trees for single-cell data of a muscle-invasive bladder tumor [26] with Scelestial, OncoNEM, SCITE, and BitPhylogeny. These data were obtained by single-cell exome sequencing of 44 tumor cells as well as exome sequencing of normal cells, and included 443 variable sites. We converted this to the required input matrix for OncoNEM, SCITE, and BitPhylogeny, in which only the reference state, variant state, and missing values were specified. In the resulting matrix, 27% of the elements represented the reference state, 17% represented variant states, and 55% represented missing values. Unlike the original study [5], we used all cancerous cells as well as 13 normal cells for lineage tree reconstruction, to see whether a distinct cancer cell lineage would become apparent in the inferred trees. All methods except SCITE removed the redundant inner nodes of the trees (i.e., nodes with not more than one child). For the SCITE method, to obtain a comparable small tree, we compressed the tree in the same manner.

In the trees inferred by BitPhylogeny, SCITE, SiFit, SASC, and SCIPhI (Fig. 4), normal and cancerous cells were not separated into distinct subtrees.

**Fig 4:**
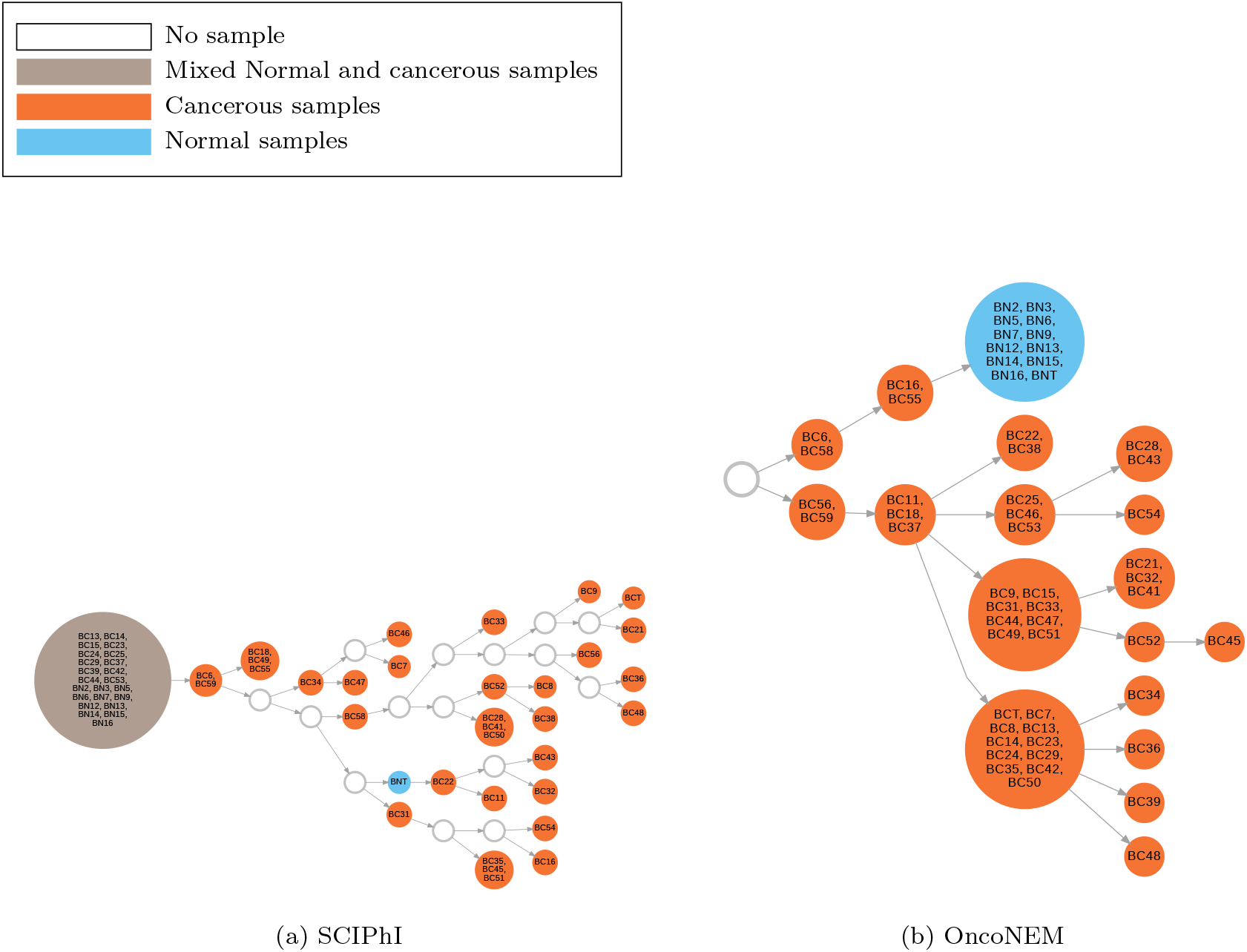
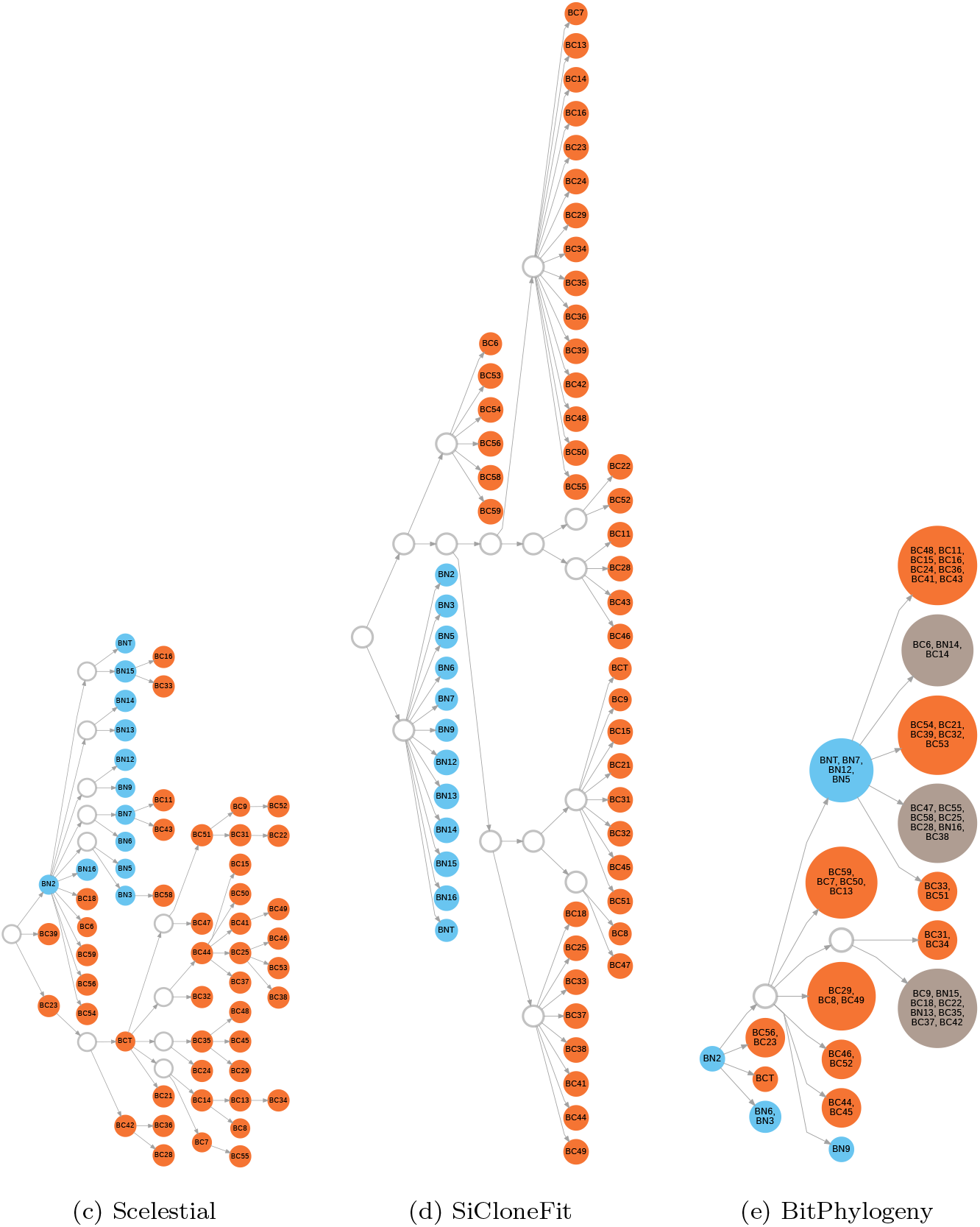
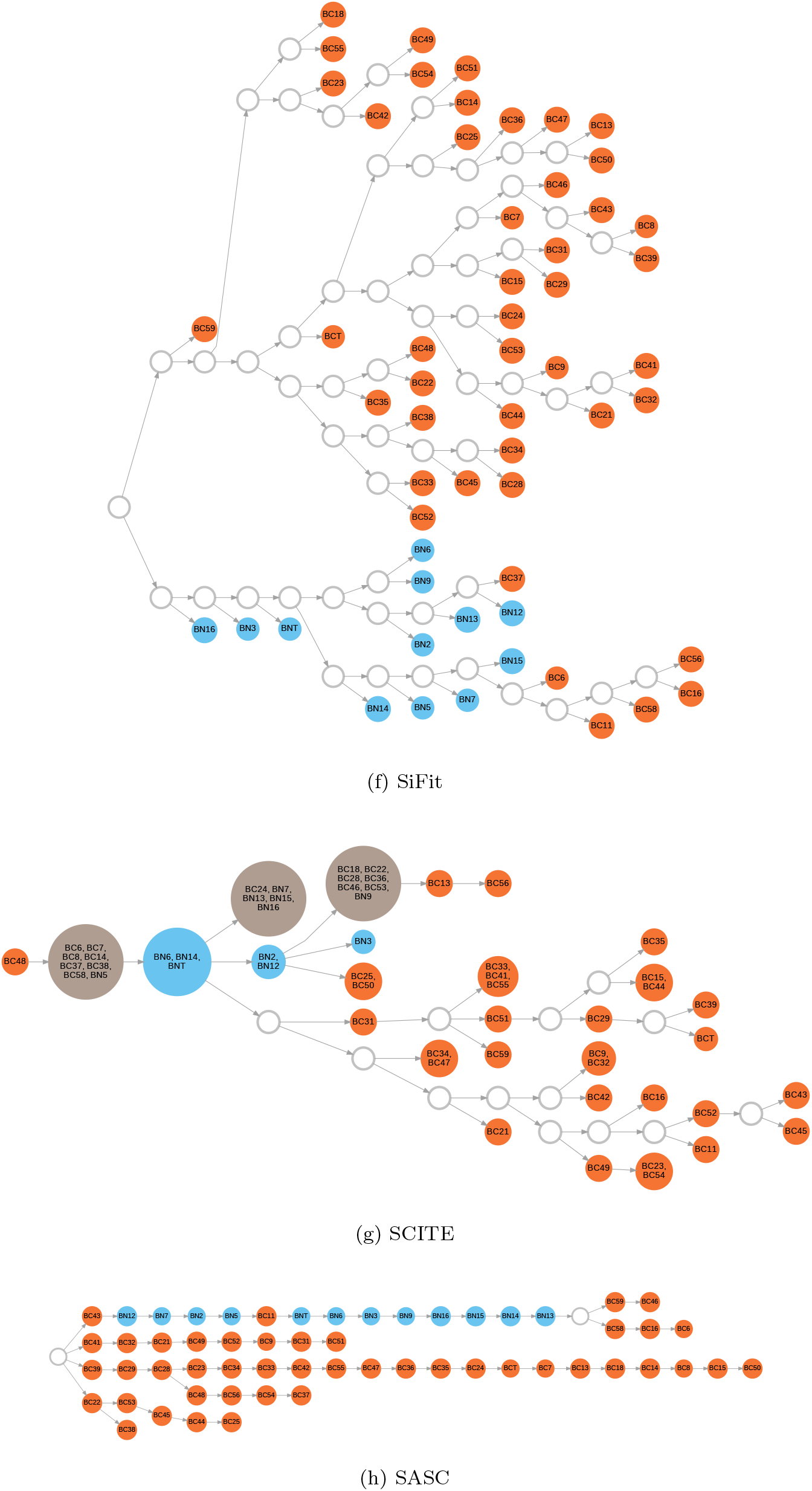
Lineage tree inferred by the different methods on a single-cell dataset from a muscle-invasive bladder tumour. Nodes in these trees represent clones (i.e., inferred, evolutionary genomes ancestral to the observed single-cell genomes). Blue nodes represent clones containing only normal cells, orange nodes represent clones containing only cancerous cells, and brown nodes represent clones containing both normal and cancerous cells. Text within the nodes indicates the identification number of the assigned sample cell(s) to the corresponding clone. White nodes represent nodes with no observed sample.

In some cases, they were even assigned to the same clone, which seems unlikely as an evolutionary scenario. In contrast, the trees of OncoNEM and Scelestial effectively separated normal and cancerous cells. OncoNEM placed all the normal cells in one clone and Scelestial separated all the cancerous cells and normal cells into distinct subtrees.

The evolutionary trees (Fig. 4) returned by all the algorithms except Scelestial were directed. The most plausible scenario for cancer evolution is the rooting of a cancer lineage close to this root or to some internal lineage of normal cells, instead of cancer cells being placed as ancestral to normal cells. OncoNEM’s tree includes a normal cell as a root, followed by some cancerous colonies placed between the descendant normal cells and this root.

### 2.4 Case study 2: a single-cell dataset from metastatic colorectal cancer

We inferred lineage trees by all methods on single-cell data of colorectal cancer from two patients [25]. In this dataset, 178 single-cell samples were gathered from the first patient: 117 samples from normal cells, 33 samples from the primary tumor, and 28 samples from metastasis of the tumor to the liver. The data represent 16 genomic sites with 6.7% missing values.

In the reconstructed lineage trees for the first patient (Fig. 5), OncoNEM, BitPhylogeny, SCIPhI, and SCITE suggest clones including both normal and cancerous cells, which, as outlined above, is unrealistic. SASC and SiFit produced lineage trees with several normal and cancerous cells that evolved from cancerous clones, while Scelestial again separated the two types of cells into distinct sub-clades: except for three metastatic cell samples, Scelestial suggested evolution of the primary tumor from normal cells and evolution of the metastatic tumor from the primary tumor. Like Scelestial, SiCloneFit placed the same three metastatic (misplaced) samples in the Scelestial tree between normal and primary tumor cells. However, the tree generated by SiCloneFit suggested independent evolution of the metastatic cells and primary tumor cells, which seems less likely, though not impossible.

**Fig 5:**
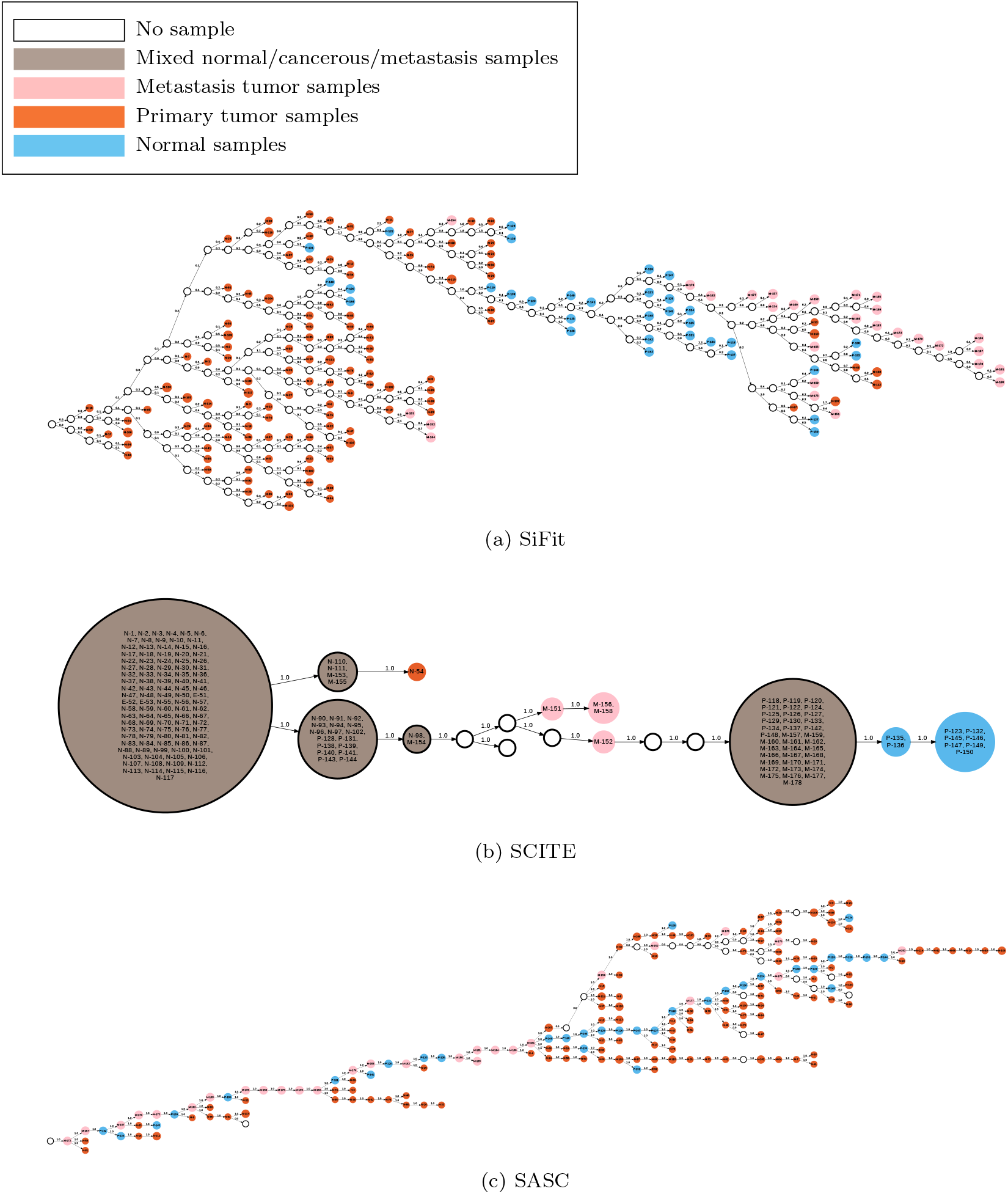
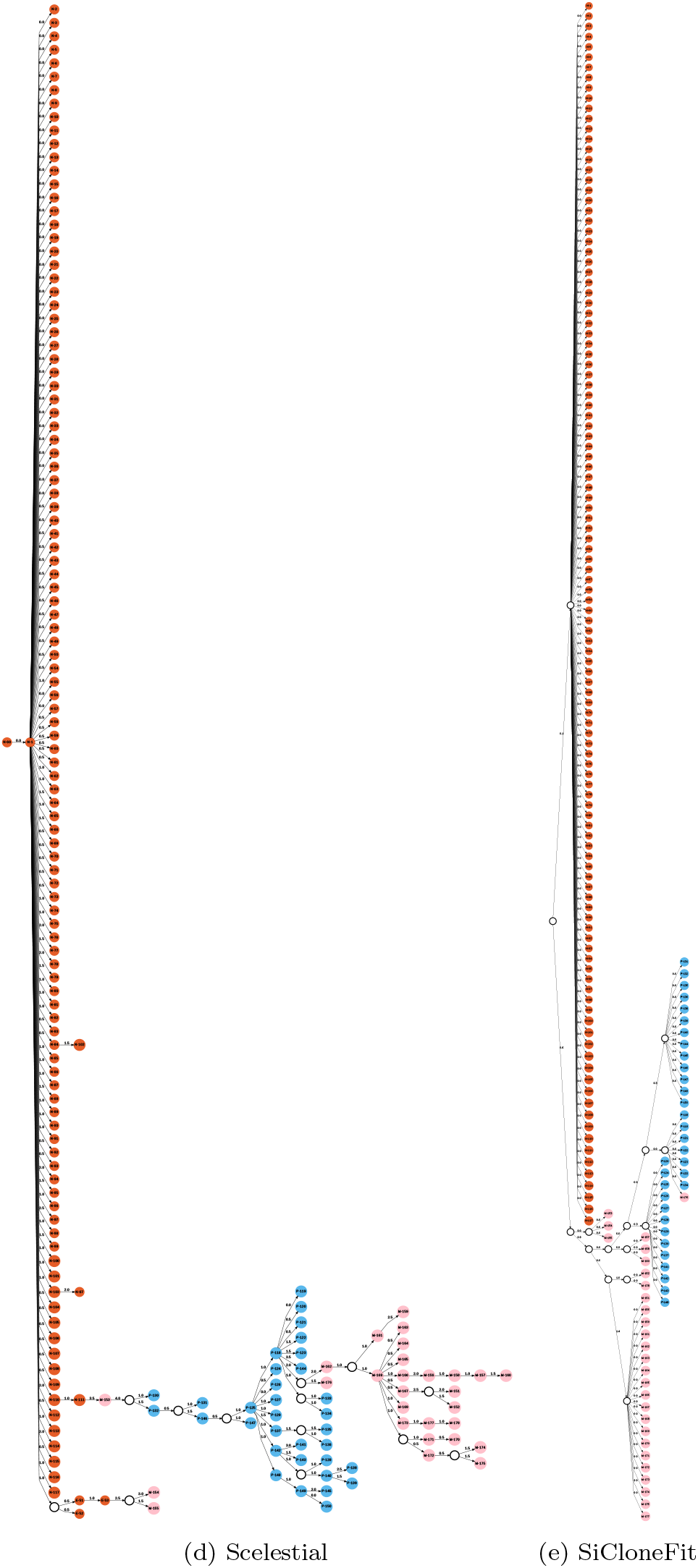
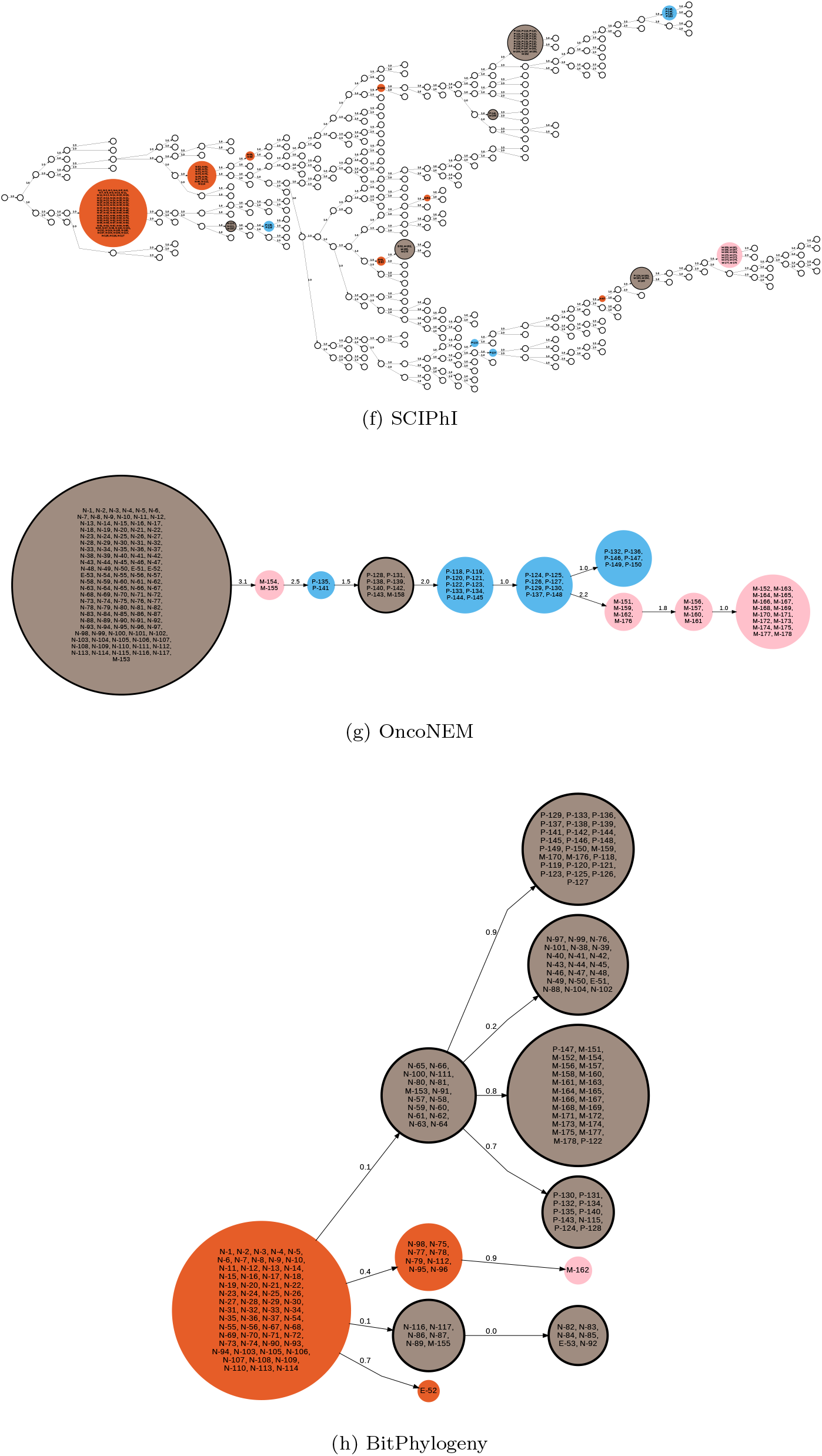
Lineage tree inferred by the different methods on a single-cell dataset from the first colorectal cancer patient.

However, SiCloneFit placed one metastatic cell (M176) closer to the primary tumor. Taken together, Scelestial performed best at separating the normal cells, primary tumor, and metastatic cells. SiCloneFit was almost as good, with only one more seemingly misplaced cell.

From the second patient, 181 single-cell samples were taken: 113 normal cells, 29 primary tumor cells, and 39 metastatic cells. Thirty-six features were extracted by evaluation of the cancer mutations of the samples. Of all sites, 7.7% were missing values.

All algorithms separated normal from cancerous cells less well than for the first patient (Fig. 6). Similar to their trees for the first patient, OncoNEM, BitPhylogeny, SCIPhI, and SCITE suggested several mixed clones including normal and cancerous cells. SASC and SiFit produced lineage trees with several normal and cancerous cells evolving from each other. In SiFit’s tree, there were fewer normal cells evolved from cancerous cells that in SASC’s tree, but instead had many distinct subtrees with cancerous cells descending from normal cells. Except for one cell (M-175), all cancerous cells were separated from normal cells into one subtree in Scelestial’s tree. In the SiCloneFit tree, the cancerous cells M-175 and M-178 were also misplaced outside of a cancer lineage and closer to normal cells. Metastatic and primary tumor cells were not well separated from one another in the trees generated by Scelestial as well as by SiCloneFit. Overall, Scelestial and SiCloneFit created the most realistic lineage trees of all the methods analyzed.

**Fig 6:**
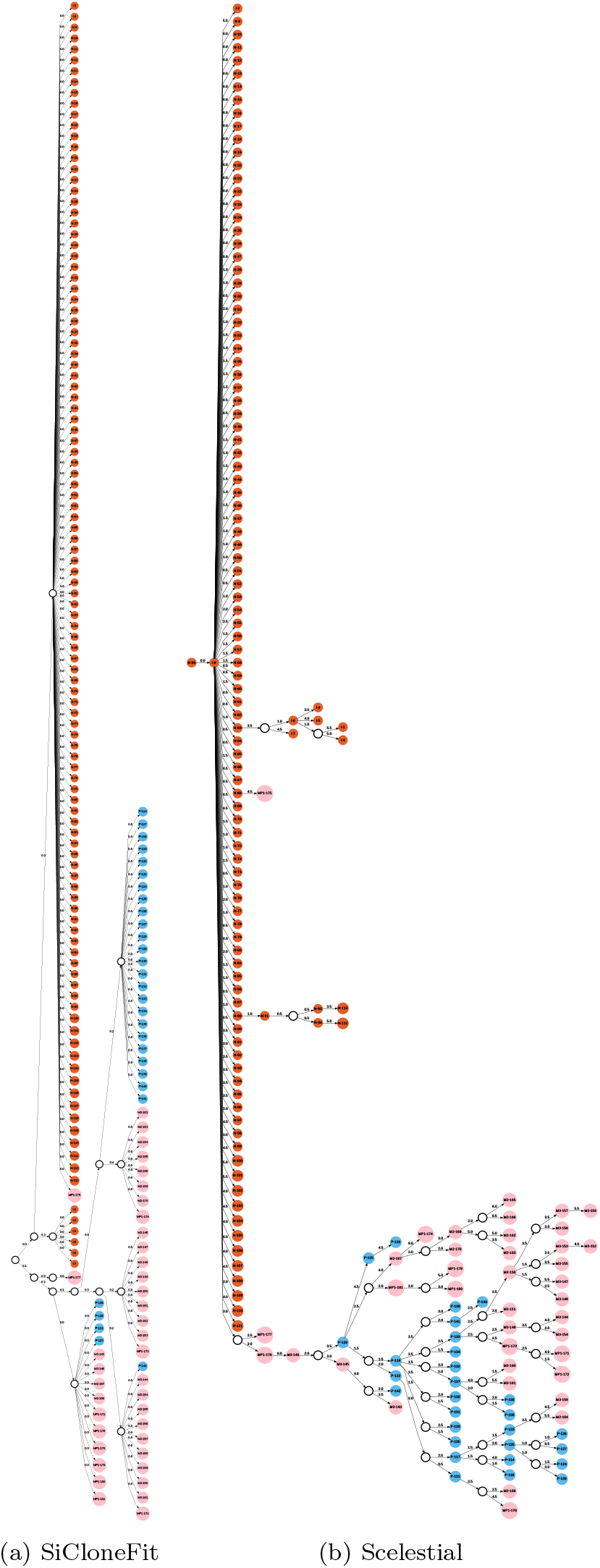
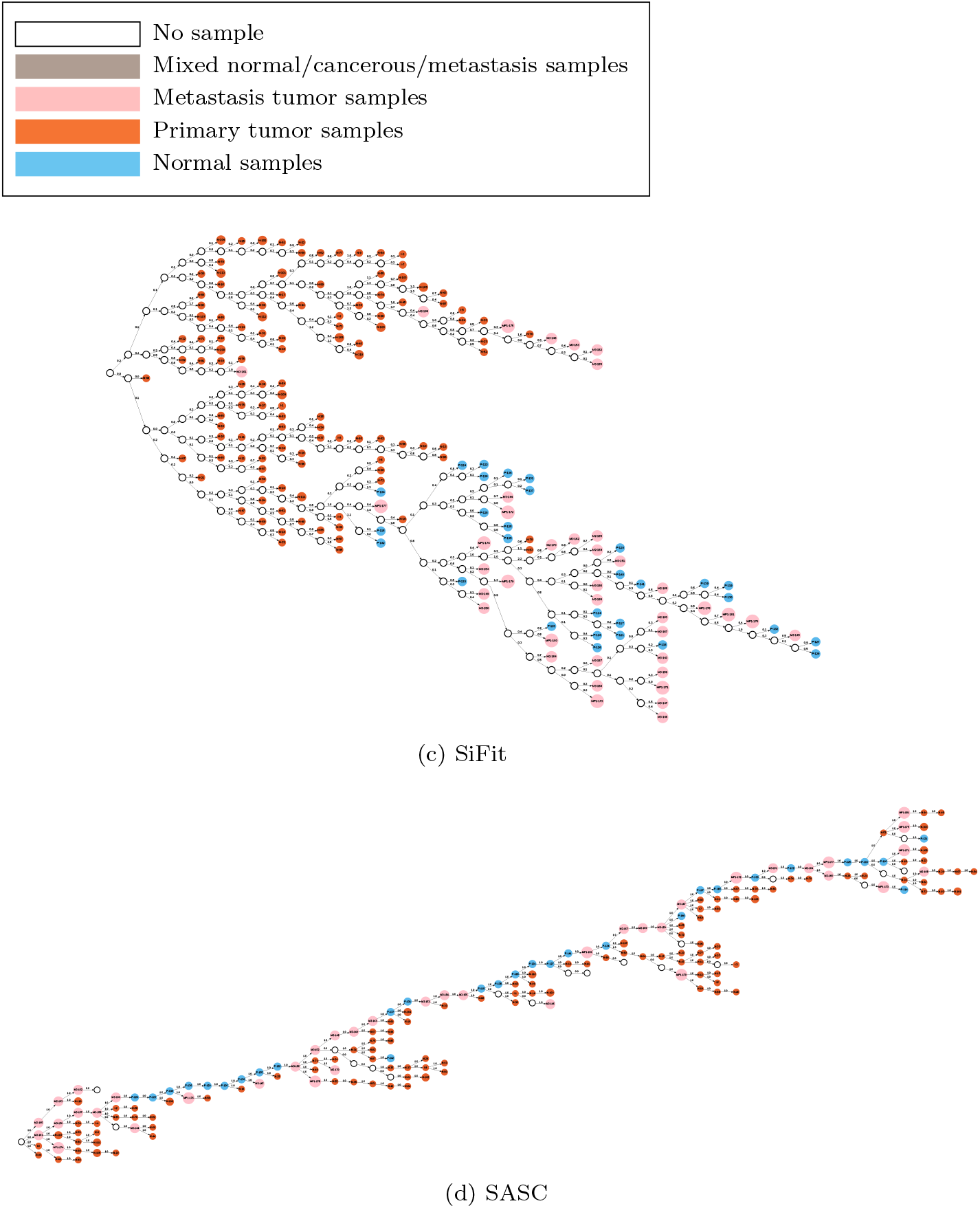
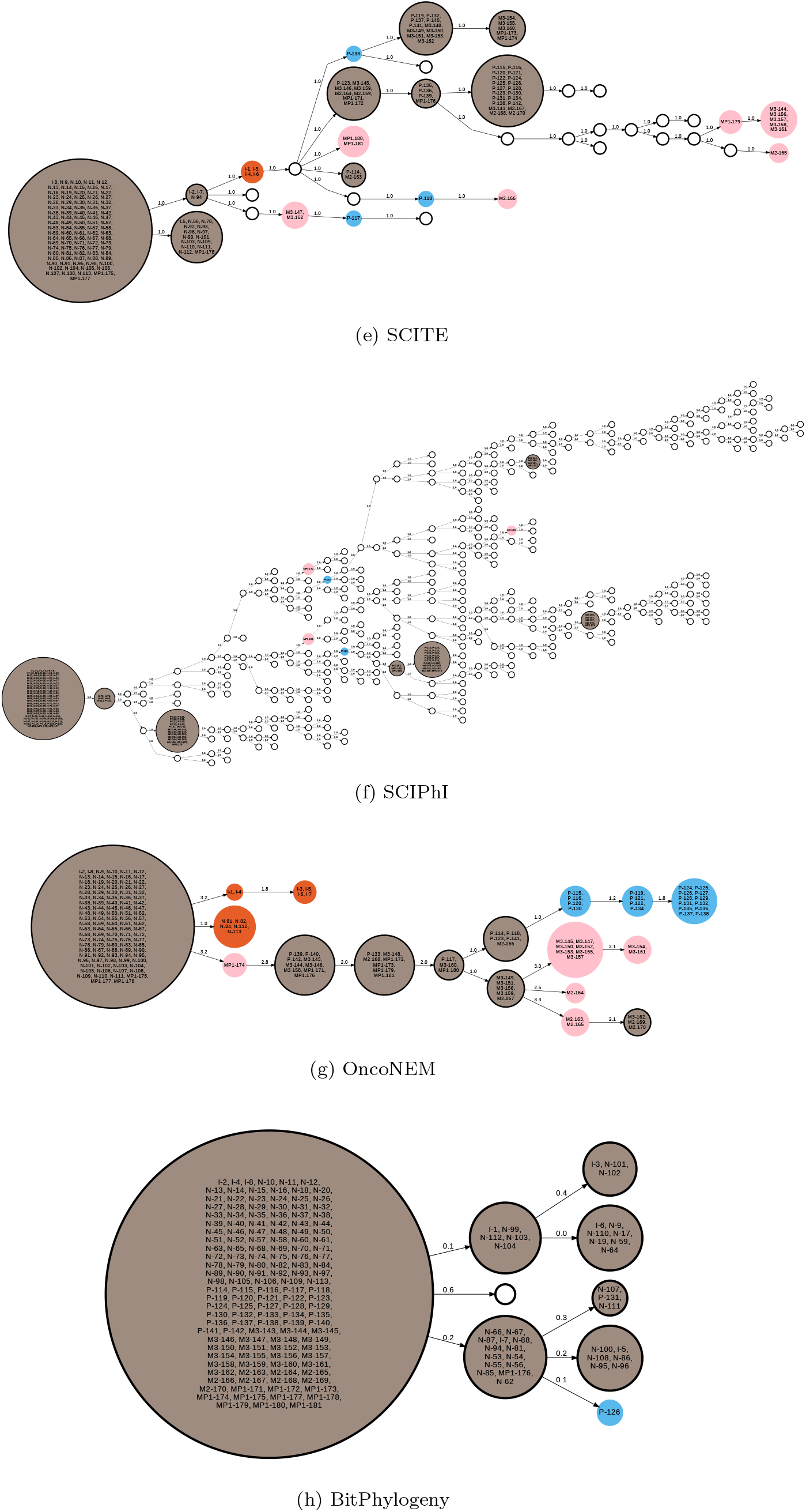
Lineage tree inferred by the different methods on a single-cell dataset from the second colorectal cancer patient.

### 2.5 Run time efficiency

We compared the run times of the eight methods with default settings on the 110 datasets generated by the OncoNEM simulator (Fig. 7). We simulated data with 10 mutations on average at each evolutionary step, a range of 20 to 100 samples, and a range of 50 to 200 sites. We only included pairs of methods and cases for which the method’s run time was less than 15 minutes.

SCITE, SCIPhI, BitPhylogeny, and Scelestial were the only methods finishing their task across the whole range of sample sizes within 100 seconds. Scelestial, SCIPhI, and SCITE’s run times grew almost linearly with an increasing number of samples. The run time of BitPhylogeny does not seem to be directly related to the number of samples but was rather random for the tested cases, since it applies MCMC and its run time only depends on the number of iterations. When we increased the number of sites, the run time of Scelestial also grew almost linearly. Next to Scelestial, BitPhylogeny and SCITE were the fastest methods. Over all cases, Scelestial was faster than all other methods in seven cases; BitPhylogeny was the fastest in six cases.

**Fig 7:**
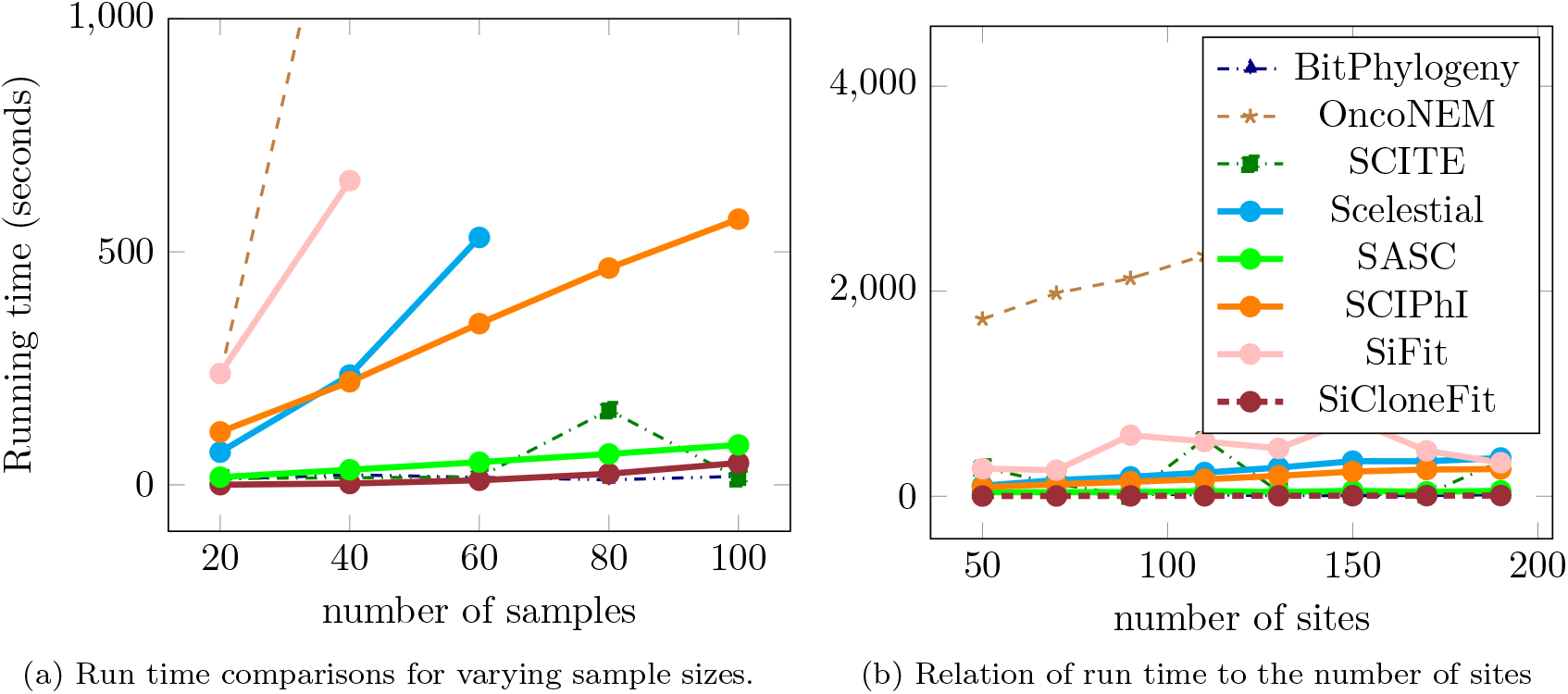
Run time comparison in relation to the number of samples and sites.

With these data, as well as being fastest, Scelestial, as before, also had the smallest average error in lineage tree reconstruction (Fig. 8b). Among 130 cases, in terms of sample distance error, Scelestial performed best in 112 cases and OncoNEM performed best in 22 cases. In terms of the split similarity measure of topological correctness, Scelestial performed best in 112 cases, OncoNEM in 18 cases, BitPhylogeny in four cases, and SCITE in two cases. In case of ties, best performance was counted for all methods. OncoNEM was slightly more accurate in some cases; however, it had a substantially larger run time. Thus, in these tests Scelestial was best overall in both run time and error. SCITE and BitPhylogeny had similar split similarity, with SCITE being faster than BitPhylogeny on average. Considering the pair distance error, the performance of BitPhylogeny was slightly better than that of SCITE. The performance of OncoNEM with respect to the pair distance error was better than that of BitPhylogeny and SCITE in a lot of cases and on average. However, the variance of the pair distance error and the split similarity of OncoNEM’s results were higher than those of others. The range of variation of performance for OncoNEM is clearly shown in the split similarity measure chart for this dataset (Fig. 8b).

**Fig 8:**
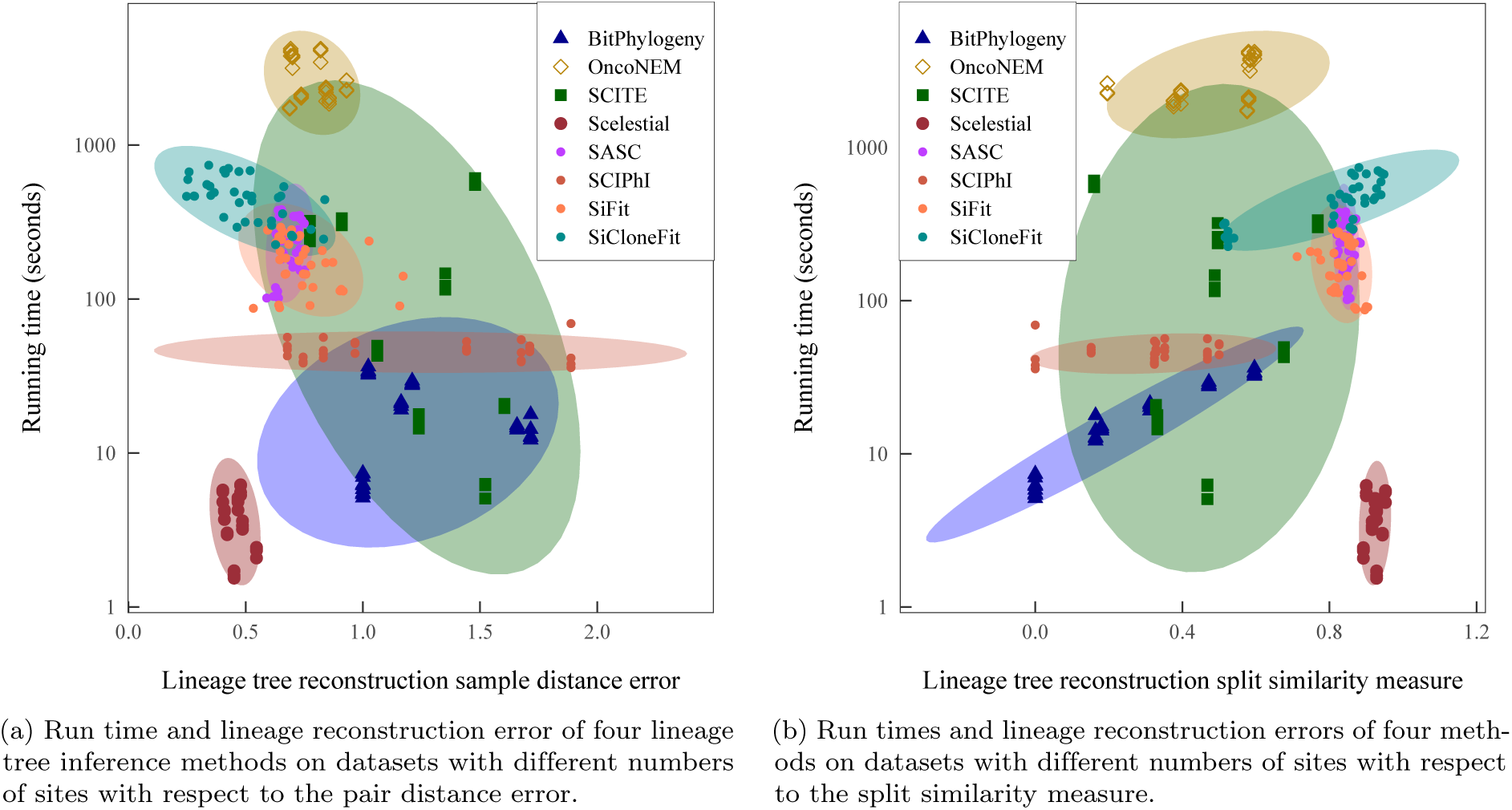
Comparison of methods with respect to running time and lineage tree reconstruction error.

According to theoretical analysis, the run time of Scelestial is polynomial with respect to the number of samples with exponent *k*, which is a parameter for the Steiner tree approximation algorithm. In Fig. 7, the quadratic form of the run time might not be directly evident. This is normal because the chart is cropped for large values to show the differences for all the algorithms except OncoNEM and SiFit. The chart is also drawn logarithmically in the y-axis, which makes it hard to see the actual growth in it. SCIPhI, SCITE, and BitPhylogeny show similar behavior. In practice, the run times of all the algorithms except for Scelestial and SCIPhI grew substantially with an increasing number of sites.

## 3 Materials and Methods

### 3.1 Data format

We model the data as an *m* by *n* matrix *D* with single-cell samples as columns and features as rows. The element of *D* corresponding to a single-cell *c* and a locus *f* is denoted *D*[*c, f*] and represents the result of a variant call obtained from a single-cell sample of a diploid genome, which may be one of the 10 states from the set {*A/A, T/T, C/C, G/G, A/T, A/C, A/G, T/C, T/G, C/G*} or a missing value *X/X*. When the data do not provide all the information (e.g., for data obtained from OncoNEM’s simulation tools), we can convert 0/1 (reference state/variant state) matrices to the 10-state format by coding 0 as A/A and 1 as C/C. All the single-cell lineage tree reconstruction methods we consider here support missing values in their input matrices.

With this coding, we cannot differentiate between the case of two similar alleles, e.g. A/A and the case of a missing allele in one strand. We hope that considering an error as a regularization method helps us to find the model (which is a lineage tree) that fits the data best.

### 3.2 Synthetic data generation via tumor simulation

We developed a tumor growth simulation method as a data source for the evaluation of Scelestial and for a comparison with state-of-the-art methods. The simulation has three phases: (1) simulation of evolution, (2) sampling from the tree, and (3) simulation of sequencing.

In the simulation of evolution phase, the evolutionary process is simulated and a tree is generated. This simulation is based on evolutionary events that happen in a tumor. We modeled the cell division, mutation, and selection in the evolutionary process of the tumor in this simulation. Each node in the evolutionary tree represents a cell (or representative of a bulk of similar cells) in the history of tumor formation. To each of these, an advantage value is assigned that shows the relative growth or division advantage of the cells corresponding to that node. The advantage value of a node is the same as its parent advantage value plus or minus a uniform random number. The evolutionary tree is constructed through several steps. In each step, one new node is generated and its parent is chosen from the current nodes with a probability proportional to their advantage values. We calculated the advantage value for the newly born node as a random perturbation added to its parent’s advantage value. The actual sequence for the new node is calculated from the sequence of the parent with some random mutations. The number of mutations from its parent is derived from a Poisson distribution about an average parameter specified in the input. The locations of the mutations are chosen uniformly from all the loci.

In the sampling phase, samples are chosen from the nodes of the tree with a probability which is proportional to their advantage values. In the simulation of the sequencing phase, missing values and errors are then incorporated by stochastic processes.

### 3.3 Synthetic data generated by OncoNEM’s simulation tool

We used simulated data generated by OncoNEM’s simulation software for our evaluation. The OncoNEM simulator is based on the evolution of clones. First, it generates a lineage tree, then it assigns mutations to tree nodes under the infinite site assumption. Afterwards, it generates output sequences by sampling from the tree and generating sequences with single-cell sequencing issues, including missing values, false positives, and false negatives.

### 3.4 Comparison of a resulting lineage tree with a ground truth

We define two measures between trees: (1) a distance measure that compares distances between samples in two trees, and (2) a similarity measure that compares the set of splits made by two trees applied to the set of samples. For both measures, we used a weighted version of classic 0/1 measures.

Different measures used in the literature are clustering accuracy [12], order of mutation in a phylogenetic tree, ancestor-descendant accuracy of mutations, different lineages, and co-clustering mutations [12]. These measures are used for evaluating mutation trees, which is not what we generate in the Scelestial method.

#### 3.4.1 Sample distance measure between trees

We define the sample distance measure between two trees based on the shortest-path matrix idea proposed by [5]. The basic idea is that we calculate pairwise distances between each pair of input cells in a tree *T*. We also create a pairwise distance matrix *PD_T_* between the inputs for the tree. Following this, we normalize the matrix *PD_T_* to obtain 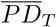 as 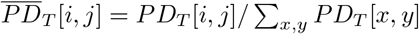. Since different concepts of weight for the tree edges are used in different methods, the normalization phase allows us to neglect absolute values and only consider the relative edge distances in the lineage trees provided. We define the distance between two trees *T* and *T*′ as 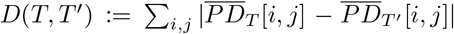, which represents the distance between the normalized pairwise distance matrices for the two trees. The value of *D*(*T, T*′) lies in the range between 0 and 2.

#### 3.4.2 Split similarity measure between trees

We defined the split similarity between two trees *T* and *T*′ as the similarity between two sets of splits generated by two trees. For each edge *e* in a tree *T*, we define the split *S_e_* as the set {*A, B*}, where *A* and *B* are the set of samples separated by the edge *e* in *T*. We define a similarity between two splits {*A*_1_*, B*_1_} and {*A*_2_*, B*_2_} as the number of samples that are split similarly in two splits. However, since there is no difference between *A* and *B* in two splits, we define the similarity score as:

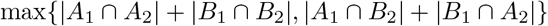

To calculate the distance between two sets of splits, we find the mapping between the elements of two sets with the maximum similarity score. The mapping similarity score is the sum of the similarity of the scores of matched splits. We than defined *D_S_*(*T, T*′) as the normalized matching score which is the mapping score between *T* and *T*′ divided by the mapping score of *T* with *T*. The split score value is between 0 and 1.

### 3.5 The Scelestial algorithm

The Scelestial algorithm is based on an established theoretical computer science problem called the Steiner tree problem. We incorporate an approximation algorithm for this problem provided by Berman et al. [24] and its modification for lineage tree reconstruction [23]. We modify the algorithm to support missing values and imputation. In the following, we examine the details of the Steiner tree problem, the reduction of lineage tree reconstruction to the Steiner tree problem and the incorporation of missing values into our method.

A schematic view of the Scelestial algorithm is illustrated in Fig. 9. The main part of the Scelestial algorithm is described in Algorithm 1. Sub-modules of the Scelestial algorithm are presented in Algorithm 2–5.

**Fig 9:**
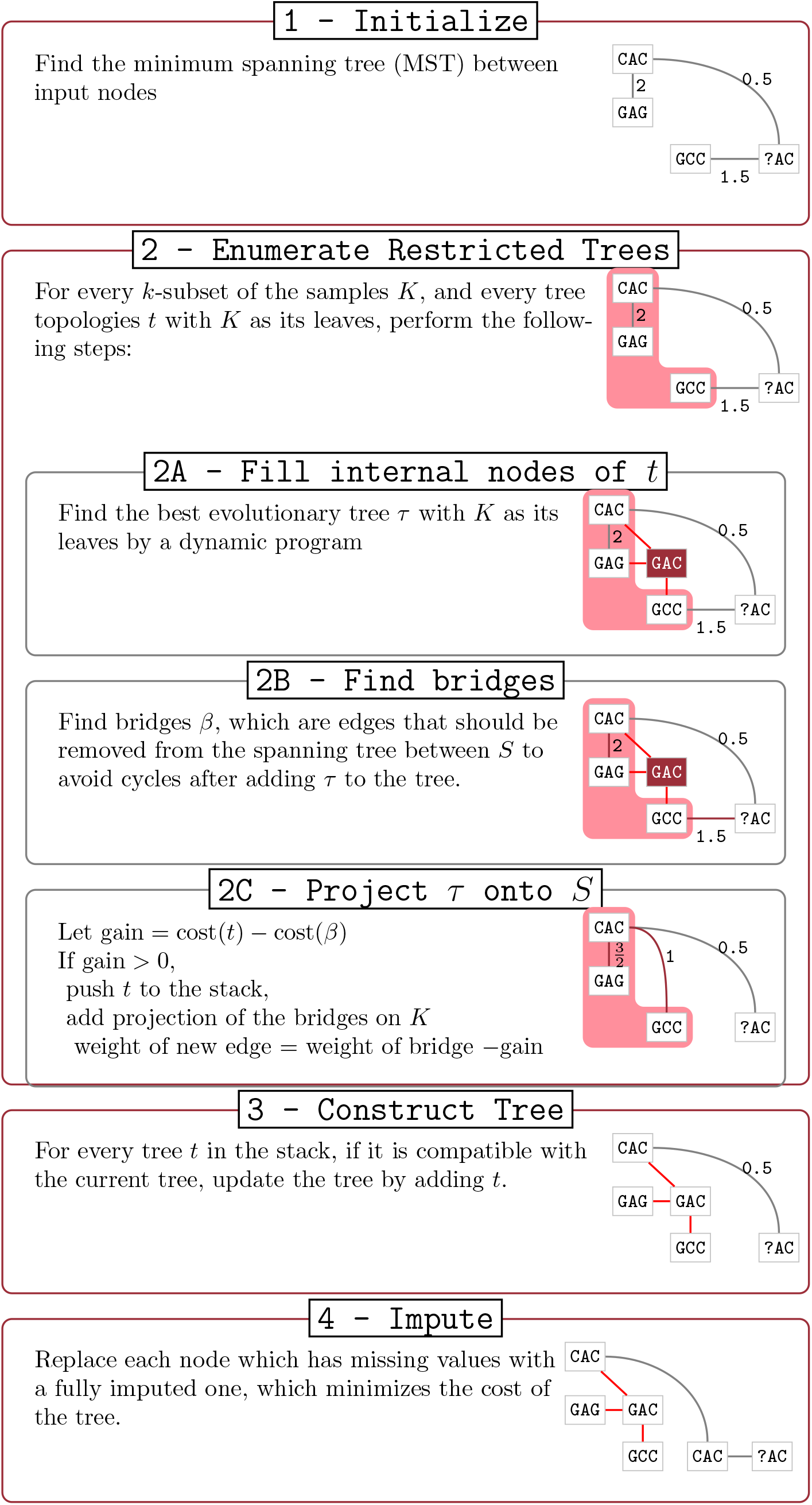
Illustration of the Scelestial algorithm.

#### 3.5.1 The Steiner tree problem

The Scelestial algorithm is based on the Berman approximation algorithm for the Steiner tree problem [24]. The input of a Steiner tree problem consists of a weighted graph *G* = (*V, E, w*) and a subset of its vertices *S* ⊆ *V*, which are called terminals. It is convenient to suppose that the weight function *w* satisfies the triangle inequality (i.e. *w*(*x, y*) + *w*(*y, z*) ≤ *w*(*x, z*), for all three vertices *x, z, y* ∈ *V*). In the case of no missing values occurring in the data, the triangle inequality constraint is satisfied, for example, for the Hamming distance between nodes. However, in the case of single-cell data, which contain a lot of missing values, we should consider this constraint carefully.

The Steiner tree problem is the problem of finding a minimum-weight connected subgraph of *G* that contains all the terminals *S*. The Steiner tree problem is an NP-hard problem and it is known that under reasonable complexity assumptions, no approximation algorithm can approximate the result better than a factor of 96/95 ≈ 1.0105 [27].

Most approximation algorithms for the Steiner tree problem focus on results that consist of a set of subtrees, each having *k* terminals at most, for some constant *k*. A solution for the Steiner tree problem with this property is called a *k*-restricted Steiner tree. We call the value *k* the restriction number of a Steiner tree. Borchers and Du [28] showed that restriction of the search space to *k*-restricted Steiner trees does not change the approximation factor of an algorithm too much. More specifically, if *k* = 2^*r*^ + *s* where 0 ≤ *s* ≤ 2^*r*^, the restriction to *k*-restricted Steiner trees reduces the approximation ratio to

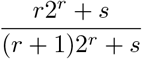

#### 3.5.2 Berman’s approximation algorithm for the Steiner tree problem

The Berman et al.’s [24] approximation algorithm consists of three phases: examination, evaluation, and application.

During the examination phase, the algorithm maintains a minimum spanning tree *M* on the terminal set *S*. In this phase, the algorithm considers all the subsets *K* of the terminals with a size of at most *k* (for some fixed constant *k*) and all the topologies *τ* for the trees with *K* as its leaves. For each *K* and *τ*, the algorithm finds the best lineage tree *t* (i.e., the best sequences for the internal nodes). This could be done through dynamic programming. By adding *t* to the spanning tree, *M* will create (*k* − 1) cycles. The algorithm finds a subset of edges of *M* with maximum cost to be removed from *M* + *t* to obtain a tree again. These edges are called bridges *β*. A value gain is defined for *t*, which is equal to the amount of decrease in the resulting tree if we decide to incorporate *t*, which is equal to cost(*β*) − cost(*t*), where the cost of a set of edges is defined as the sum of the costs of its elements. If the the gain is greater than 1, the algorithm adds *t* to a stack for the evaluation phase, it removes *β* from *M*, and adds some new edges to *M* instead. For each edge *e* of *β*, the algorithm finds the vertices *u* and *v* from *K* which are going to be disconnected after removing *e*. It adds the edge (*v, u*) to *M* with the cost cost(*e*) − gain. At the end of the examination phase, we will have a stack of some trees and *M*.

In the evaluation phase, the algorithm pops the trees *t* one by one from the stack. It starts with an initially empty set of edges *M*′. At each step it checks if *t* does not make a loop with *M*′. If this is the case, the algorithm accepts *t*; otherwise, it rejects *t*. Finally, in the application phase, the algorithm merges the accepted trees.

The run time of the algorithm for a general Steiner tree problem is 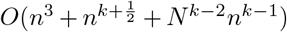, where *N* is the number of vertices in the graph. Note that *n* is the number of terminal vertices *S*. The algorithm is an 11/6-approximation algorithm for the Steiner tree problem, if we set *k* = 3. If we set *k* = 4, the algorithm would be a 16/9-approximation algorithm. For larger values of *k*, we do not know a better approximation factor for the result of the algorithm, but the results of the execution of the algorithm show that the performance of the algorithm gets better as we increase the value of *k* [24].

#### 3.5.3 Modeling lineage tree reconstruction as a Steiner tree problem

The single-cell lineage tree inference problem can be modeled in the following way. Let *g_i_* for 1 ≤ *i* ≤ *n* be the set of input sequences over some alphabet Σ + *μ*, where *μ* is a special character representing the missing value. Each location on a sequence could be considered as a feature, which may be a locus in the sequenced genomes or may represent a genomic aberration. We suppose that all the sequences have the same length *m*. A cost function is defined for any potential lineage trees to show how well a lineage tree is fitted to the data. Normally, the cost function assigns a cost to each edge of the tree, and the cost of a tree is the sum of its edge costs. The lineage tree inference problem is the problem of finding a minimal cost lineage trees that contain all the input sequences *g_i_* and potentially some other nodes. In the resulting tree, all non-input nodes are assigned a label of length *m* from the alphabet Σ.

A general lineage tree inference problem that does not incorporate missing values could be modeled as a Steiner tree problem as follows. Let *G* = (*V, E, w*) be the graph containing all sequences of length *m* from the alphabet Σ that have edge weights derived from the tree cost function. Thus, the lineage tree problem would be equivalent to finding a Steiner tree in this graph with the input sequences as its terminal set *S*.

#### 3.5.4 Incorporating an approximation algorithm for lineage tree reconstruction

We showed how lineage tree reconstruction could be modeled as a Steiner tree problem in the previous section. However, through this method, the size of the graph is |Σ|^*m*^, which is exponential to the length of input of the lineage tree reconstruction problem. On the other hand, some approximation algorithms do not consider all the vertices of graph *G* and they only work with the best Steiner trees over certain subsets of terminals. Since we can solve the Steiner tree problem over a small subset of terminals, we can use these approximation approaches.

To apply Berman et al.’s algorithm for lineage tree reconstruction, as already suggested by Alon et al. [23], for every subset *S*′ ⊆ *S*, with a of size at most *k*, we consider all the possible tree topologies with *k* leaves at most. Since the number *k* is constant, the number of these tree topologies would also be constant. Next, for the pair consisting of a subset *S*′ and a tree *T*, we find the minimum-cost Steiner tree by dynamic programming. The rest is done by the Berman [24] algorithm. This approach gives us an 11/6-approximation algorithm for the lineage tree reconstruction problem with *k* = 3 and a 16/9-approximation algorithm for *k* = 4. The cost of an edge between two nodes with the sequences *g_i_*[*t*] and *g_j_*[*t*] for 1 ≤ *t* ≤ *m* is Σ*_t_ c*(*g_i_*[*t*]*, g_j_*[*t*]), for some cost function *c*. In this work, we assign a cost of 1 to every mutation and a cost of zero to non-mutated locations (i.e., *c*(*x, x*) = 0 for *x* ∈ Σ and *c*(*x, y*) = 1 for *x* ≠ *y* ∈ Σ).

#### 3.5.5 Support for missing values

To adapt the method to the single-cell setting, we incorporate missing values into the algorithm. We can simply do this by extending the domain of the character cost function *c* : Σ × Σ → ℝ* to *c* : Σ + *μ* × Σ + *μ* → ℝ*. To do so, we should define *c*(*μ, x*) = *c*(*x, μ*) for *x* ∈ Σ as well as *c*(*μ, μ*); *c*(*μ, x*) could be considered as the cost of imputation for a locus. One may consider the assignment of *c*(*μ, x*) = *c*(*μ, μ*) = 0. However, this assignment violates the triangle inequality for the graph *G*. To preserve the triangle inequality between the edges’ weights, we may assign *c*(*μ, x*) = 0.5 + *ϵ* for some small constant *ϵ* (e.g., 10^−5^). Furthermore, letting *c*(*μ, μ*) = 0 does not violate the triangle inequality any longer.

#### 3.5.6 Missing value imputation

The original approach of representing lineage tree reconstruction as a Steiner tree problem does not consider the case of missing values in the input sequences. However, for single-cell data, this is important. We therefore designed a dynamic program that finds internal node sequences with non-missing values. As a result, missing values may occur only in input sequences. The translation of the missing value imputation task in our model would be the task of filling missing values with some characters from the alphabet Σ to minimize the cost function. To impute missing values after finding the lineage tree, we replace each missing value in a node with the most abundant character found within its neighbors.

**Algorithm 1.**
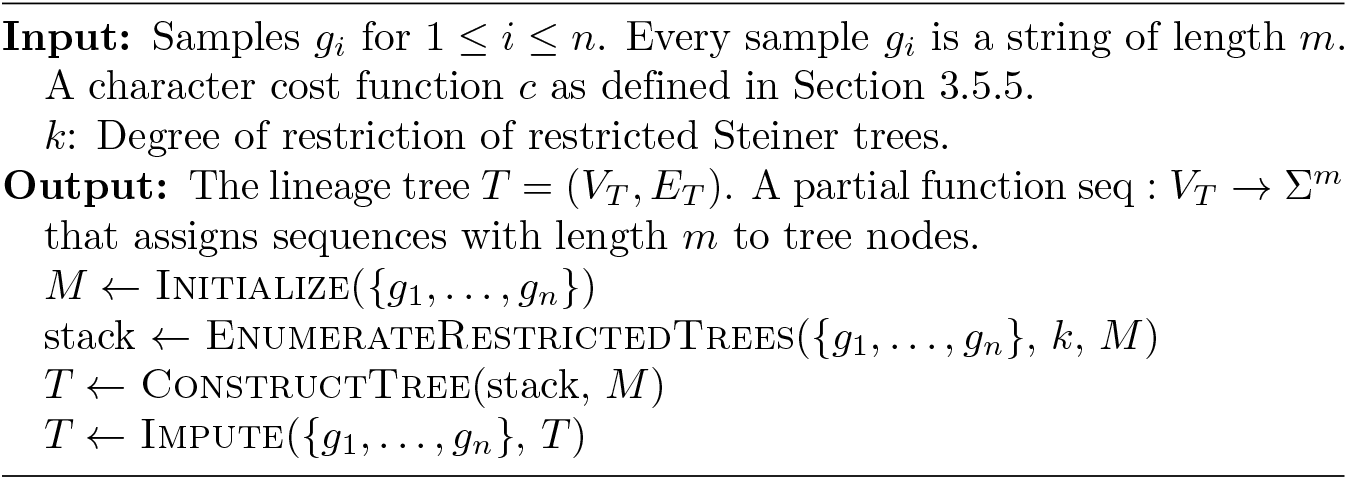
Pseudocode of the main part of the Scelestial algorithm

**Algorithm 2.**
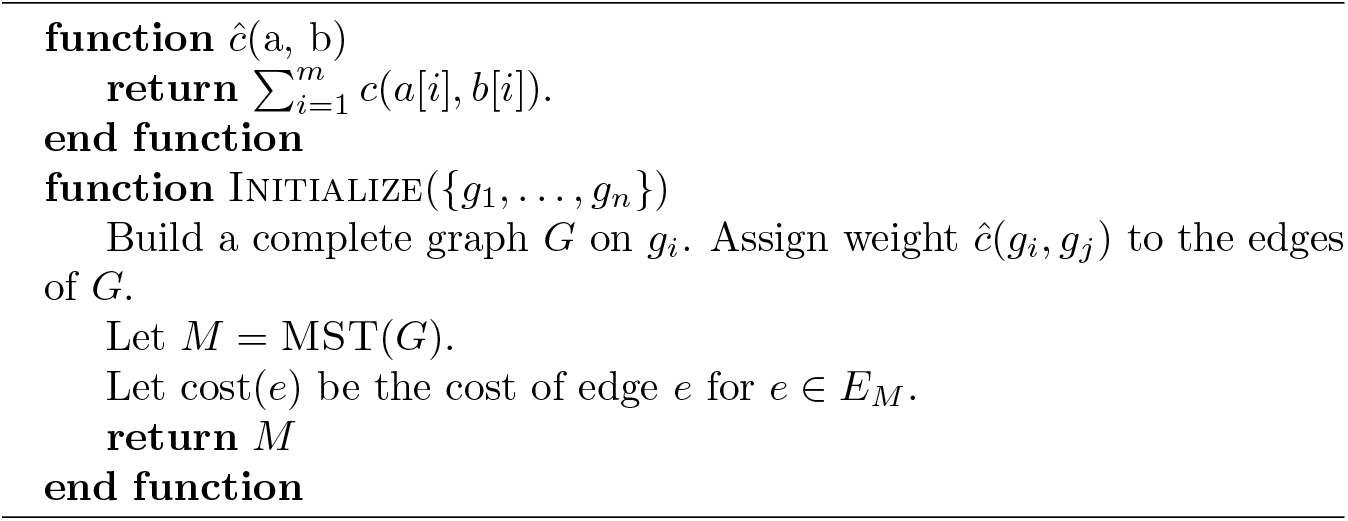
Initialize function (part of the Scelestial algorithm)

**Algorithm 3.**
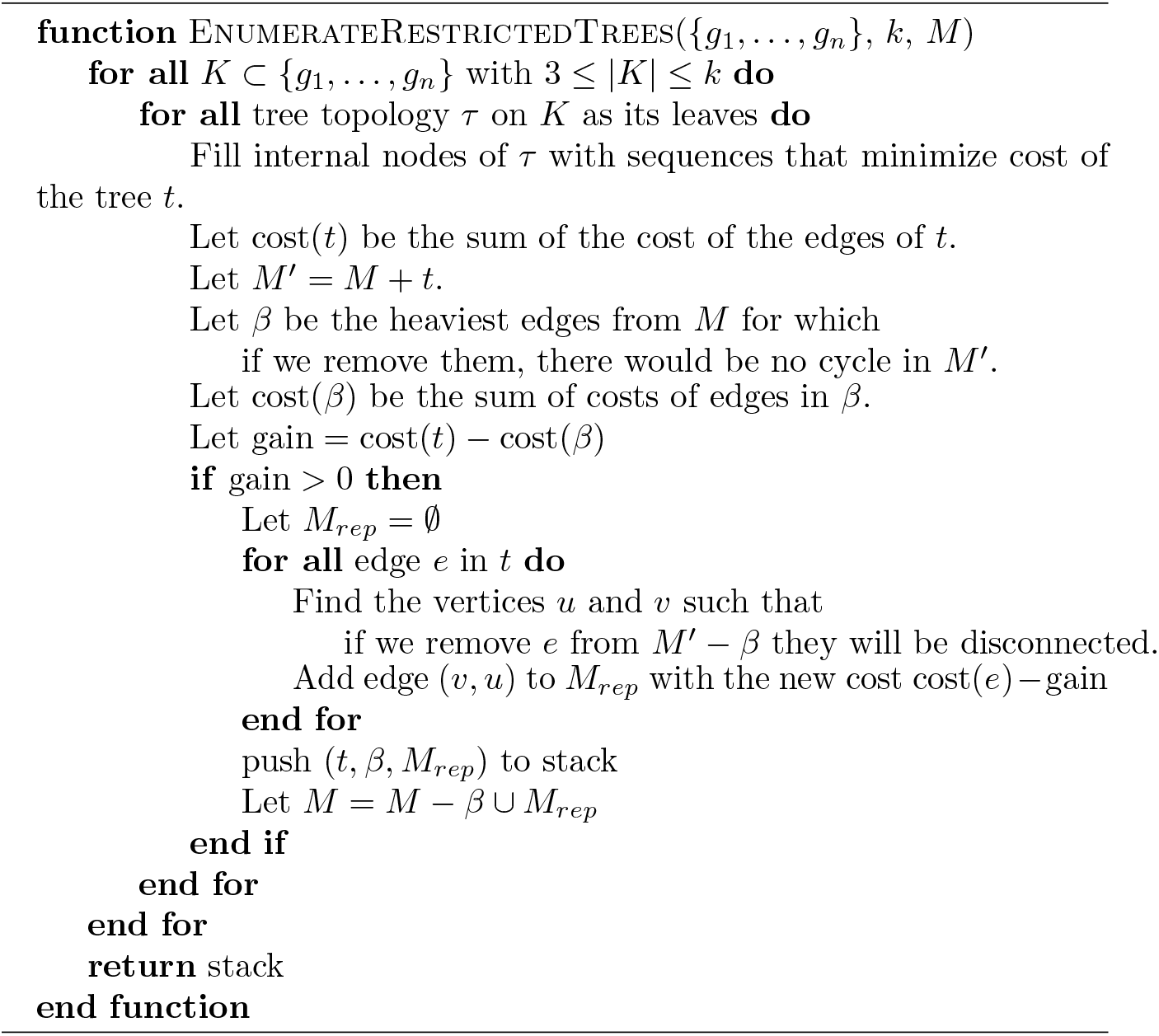
EnumerateRestrictedTrees function (part of the Scelestial algorithm)

**Algorithm 4.**
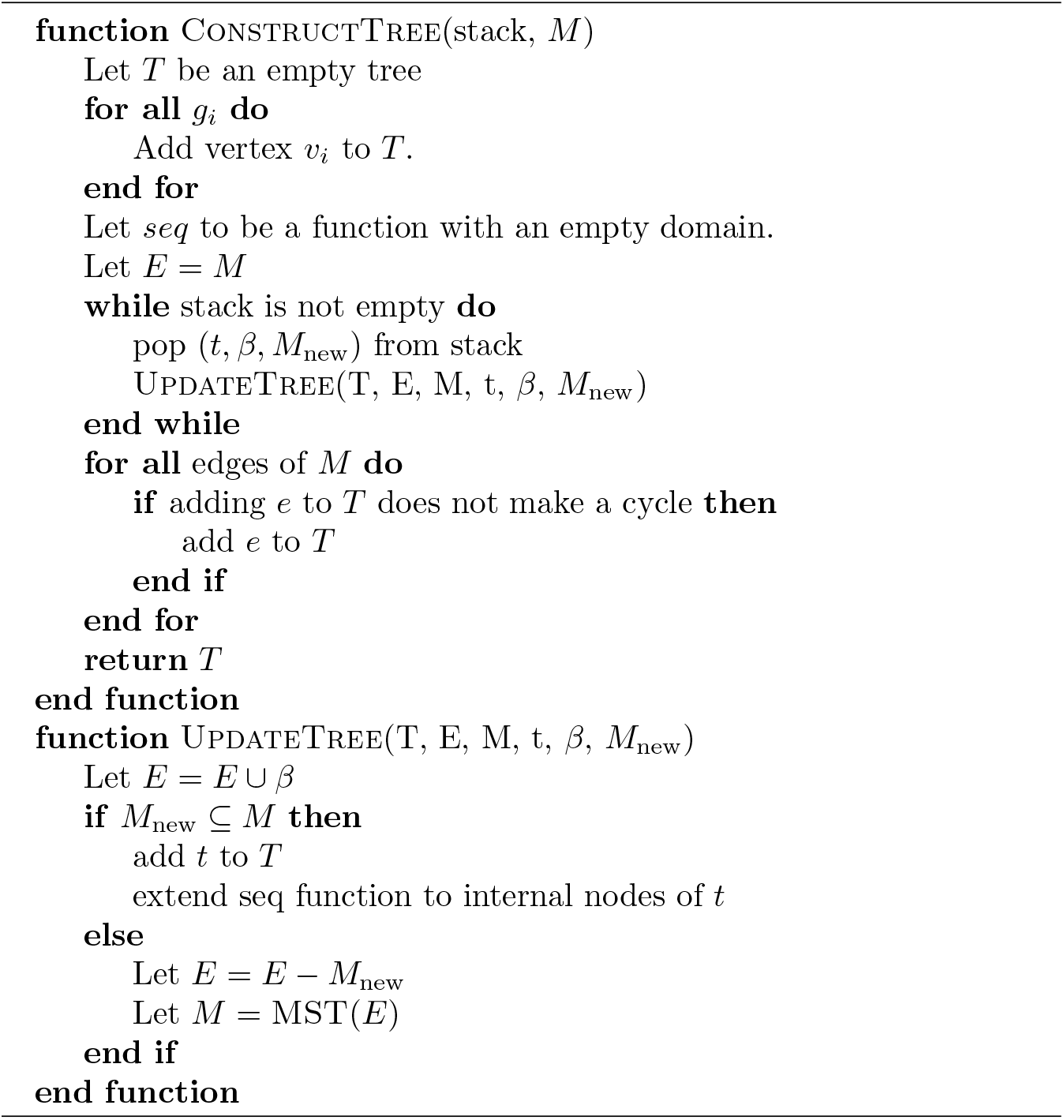
ConstructTree function (part of the Scelestial algorithm)

**Algorithm 5.**
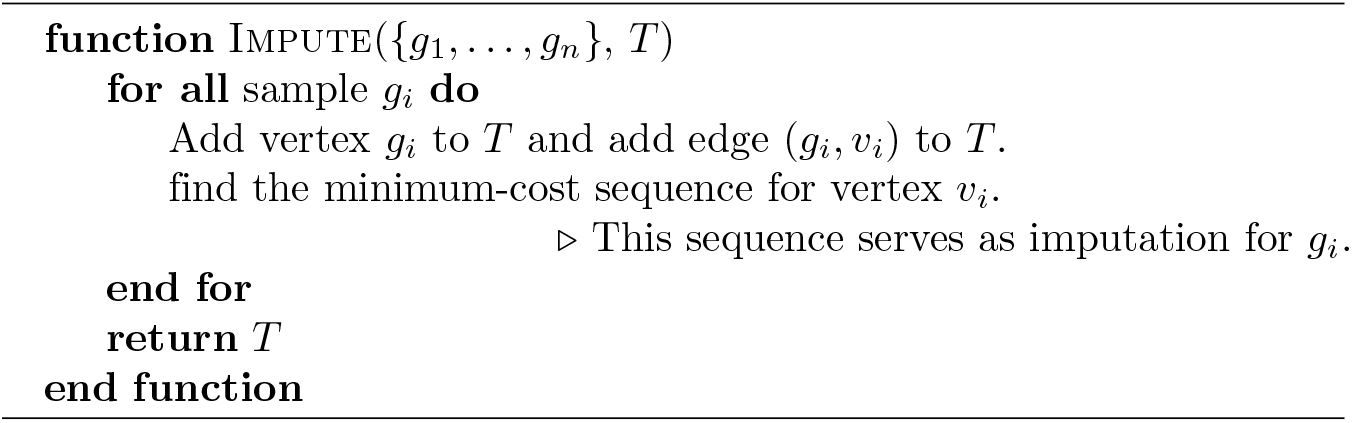
Impute function (part of the Scelestial algorithm)

## 4 Discussion and conclusions

Inference of a tumor’s evolutionary history is a crucial step towards understanding of the common patterns of cancer evolution. There are a range of methods available for phylogenetic inference with single-cell datasets. For optimal performance, many of these probabilistic approaches require substantial time for optimization (e.g., using MCMC). The increasing sample sizes of single-cell datasets emphasizes the need for fast and scalable methods in this domain that also maintain high accuracy [29].

Here, we describe Scelestial, a computationally lightweight and accurate method for lineage tree reconstruction from single-cell variant calls. The method is based on a Steiner tree algorithm with a known approximation guarantee, which we adapted to lineage tree reconstruction for single-cell data with missing values by using a variation of the Steiner tree problem called the group Steiner tree. This problem is a special case of the group Steiner problem, in which the groups are sub-hypercubes of the original graph. To achieve a fast algorithm, we further adapted the group Steiner tree algorithm through modeling a hypercube corresponding to the missing values with one representative vertex. To facilitate the use of Scelestial, we provide an implementation as a freely available R package. From another point of view, Scelestial could be considered as a generalization of the neighbor-joining method. At each step, the neighbor-joining method finds the two most similar elements (samples) and merges them to build a tree. A generalization of this idea may consider more than two samples for merging in each step. Determining a good objective function for ranking three sample candidates is not trivial. Scelestial, based on Berman’s algorithm [24], provides a generalized guaranteed neighbor-joining method.

We evaluated Scelestial’s performance with a diverse set of test cases similar to modern single-cell datasets. The datasets were generated with various ranges of clones, false positives, false negatives, and missing values, all of which were derived from real single-cell datasets [25,30]. The simulated data were produced by a tumor simulator emulating the process of tumor growth via mutation and proliferation. In this way, data for different tumors with various parameters were generated to assess computational methods over a wide range of data types.

On these benchmark datasets, we compared Scelestial with seven other state of the art phylogeny reconstruction methods, namely BitPhylogeny [4], OncoNEM [5], SCITE [6], SASC [8], SCIPhI [10], SiFit [7], and SiCloneFit [11]. As the comparisons were not straightforward, we excluded B-SCITE [12], which uses a combination of single-cell and bulk data, as well as the approach of Kim and Simons [3], where the input is hard-coded, and PhISCS [13], as it does not infer a lineage tree. Of the methods using the k-Dollo assumption, namely SASC [8], which is based on simulated annealing, and SPhyR [9], which is based on integer programming, we included SASC in the comparison. For the assessment of lineage tree quality, we applied two commonly used metrics from population genetics and phylogenetics. In this comparison, Scelestial performed best at reconstructing the ground truth tree’s topology and also the similarity of the inferred to the ground truth tree when branch lengths were considered.

When the methods were applied to cancer data, only Scelestial inferred lineage trees that separated all cancerous cells from normal cells for all three datasets, except for one cell. SiCloneFit had the second-best accuracy with two misplaced cells, and OncoNEM was the only other method that did not mix cancerous and normal cells in one single clone, although it mixed cancerous and normal cells in the evolutionary tree. The run time analysis showed Scelestial to perform up to two orders of magnitude faster than the other methods when the default settings were used. This is particularly important, as the number of single-cell genomes published in individual studies continues to be on the rise, as are multi-dataset meta-studies, as in [1].

Overall, Scelestial substantially improves lineage tree reconstruction from single-cell variant calls across key criteria such as the scalability, run times, and accuracy of lineage tree reconstruction on both real and simulated data. Cell lineage reconstructions based on diverse tumor datasets in combination with massive data gathering, resulting from fast advancing technologies, provide a better understanding of the evolutionary landscape and the associated mutations of tumors, as may also indicate the dependencies between them [31–33]. These factors may make Scelestial instrumental in furthering our understanding of the mutational landscape and the mechanisms of cancer formation and survival, as omics technologies continue to thrive. Furthermore, the results of this paper can be seen as a case study for translating concepts from theoretical computer science into advances in computational biology.

## Acknowledgments

This work was funded by a Helmholtz Incubator grant (Sparse2Big ZT-I-0007).

## Notes

### Competing Interest Statement

The authors have declared no competing interest.

